# *Bifidobacterium infantis*-mediated herpes simplex virus-TK/ganciclovir treatment inhibits cancer metastasis

**DOI:** 10.1101/2022.07.11.499524

**Authors:** Changdong Wang, Yanxi Shen, Jie Xu, Yongping Ma

**Affiliations:** Department of Biochemistry & Molecular Biology, Molecular Medicine & Cancer Research Center, College of Basic Medicine, Chongqing Medical University, Chongqing 400016, China

**Keywords:** *Bifidobacterium*, thymidine kinase, ganciclovir, tumor metastasis, gastric cancer, proteomic, bioinformatics

## Abstract

Previous studies have found that *Bifidobacterium infantis*-mediated herpes simplex virus-TK/ganciclovir (BF-TK/GCV) reduces the expression of VEGF and CD146 which implies tumor metastasis inhibition. However, the mechanism of BF-TK/GCV inhibits tumor metastasis is still not fully studied. Here, we comprehensively identified and quantified protein expression profiling for the first time in gastric cancer (GC) cells MKN-45 upon BF-TK/GCV treatment using quantitative proteomics. A total of 159 and 72 differential expression proteins (DEPs) were significantly changed in BF-TK/GCV / BF-TK and BF-TK/GCV / BF/GCV groups. Kyoto encyclopedia of genes and genomes (KEGG) pathway analysis showed some enriched metastasis-related pathways such as gap junction and cell adhesion molecules pathways. Moreover, transwell assay proved that BF-TK/GCV inhibited the invasion and migration of tumor cells. Furthermore, immunohistochemistry (IHC) demonstrated that BF-TK/GCV reduced the expression of HIF-1A, MTOR, NF-κB1-p105, VCAM1, CEBPB and CXCL12, which were associated with tumor metastasis.

In summary, besides apoptosis, BF-TK/GCV also inhibited tumor metastasis, which deepened and expanded the understanding of BF-TK/GCV anti-tumor mechanisms.

## Introduction

Tumor metastasis is the main cause of poor comprehensive treatment and poor survival prognosis(Ganesh & Massague, 2021). Despite recent therapeutic advances in cancer treatment, a large number of data show that more than 90% of tumor patients die of distal target organ metastasis. Unlike primary tumors, which can often be cured using local surgery or radiation, metastatic cancers are largely non-curable for circulating tumor cell (Chaffer & Weinberg, 2011; Ganesh & Massague, 2021; Stoletov et al, 2020). Therefore, inhibition of tumor metastasis is still a challenge in current anti-tumor research.

For high relapse rate after radical surgery and conventional therapies (radiotherapy, chemotherapy and immunotherapy) (Mun et al, 2018), novel approaches such as cancer gene therapy have raised hope to significantly improve the survival rate of tumor patients (Kenarkoohi et al, 2020; Kumar et al, 2016; Oraee-Yazdani et al, 2021; Xiao et al, 2019). Several gene therapy products have been successfully approved for clinic use (Oh & Ishikawa, 2019). Suicide gene therapy is the most frequently used method for solid tumors, which is used to eliminate tumor cells in three ways: direct killing effect, bystander effect and cell signaling pathway (Konieczny et al, 2017). Herpes simplex virus thymidine kinase gene combined with ganciclovir (TK/GCV) is one of the most in-depth-studied systems (Li et al, 2021a; Li et al, 2017; Mu et al, 2020; Pastorakova et al, 2020; Staquicini et al, 2020; Tanaka et al, 2017; Tang et al, 2009; Wang et al, 2016; Yin et al, 2013; Zeng et al, 2012). In this system, viral TK is expressed and subsequently metabolized the prodrug GCV to mono-phosphorylated GCV, which will be further converted into GCV triphosphate GCV, an analogue of deoxyguanosine triphosphate. Consequently, triphosphate GCV inhibits DNA synthesis and leads tumor cell death (Li et al, 2019b; Tang et al, 2009; Wang et al, 2016; Yanagisawa et al, 2021).

A safe, effective and controllable gene delivery system is important for gene therapy. There are three main types of vectors: viral vectors (adenoviral, adeno-associated viral), non-viral vectors (polymers, liposomes) and bacteria vectors (*Escherichia coli*, *Salmonella* and *Bifidobacterium*) (Akbulut, 2020; Guo & Huang, 2012; Zhang et al, 2013). Previous studies have demonstrated that some obligate anaerobes (*Bifidobacterium*, *Clostridium*) or facultative anaerobes (*Salmonella*, *Escherichia coli*, *Listeria monocytogenes*) can selectively accumulate and proliferate within tumors and suppress tumor growth (Duong et al, 2019; Liang et al, 2019). However, fundamental issues such as safety and targeting remain to be resolved before engineered *Salmonella* can be used in the clinic (Liang et al, 2019). *S. typhimurium* mutant VNP20009 and its derivative strain TAPET-CD had even been applied in human clinical trials (Nemunaitis et al, 2003; Toso et al, 2002). However, a significant number of bacteria can still infect normal tissue. *Bifidobacterium* is a obligate anaerobe that selectively localizes and proliferates within hypoxic regions of tumors as a non-pathogenic bacterium, and it has been considered to be an alternative strategy in tumor therapy for its acknowledge recognized safety (Wang et al, 2017; Wei et al, 2016; Xu et al, 2007; Yazawa et al, 2000; Yin et al, 2012; Zhou et al, 2016; Zhu et al, 2011). As to *Listeria monocytogenes*, it has been used to develop tumor vaccine as an intracellular pathogen.

Our previous studies have showed that *Bifidobacterium infantis* (BF)-TK/GCV inhibited the growth of gastric cancer (GC) and other several cancer models through activating extrinsic and intrinsic apoptosis pathways (Jiang et al, 2013; Jiang et al, 2014; Tang et al, 2009; Wang et al, 2016; Xiao et al, 2014; Yin et al, 2013). In addition, we detected that the expression of vascular endothelial growth factor (VEGF) and CD146 were significantly reduced after BF-TK/GCV treatment (Jiang et al, 2013; Zhou et al, 2016). It has been reported that VEGF and CD146 contributed to tumor metastasis and invasion (Ghoroghi et al, 2021; Song et al, 2021). However, it remains unclear how many molecules are involved in inhibition of tumor cell metastasis after BF-TK/GCV treatment. This study was to decipher the mechanisms of BF-TK/GCV on the inhibition of tumor cell metastasis in MKN-45 cells.

GC is one of the most common malignant tumors and the fourth leading cause of cancer death worldwide (Sung et al, 2021). Despite the gradually declining mortality, GC still burdens many countries in East Asia (Ferlay et al, 2019). Therefore, we comprehensively identified and quantified protein expression profiling in MKN-45 cells by BF-TK/GCV treatment using tandem mass tags (TMTs)-based quantitative proteomics and verified the expression of metastasis-related proteins by immunohistochemistry (IHC) in this study.

## Results

### Identification of differentially expressed proteins (DEPs)

In order to better understand the inhibition of tumor cell metastasis mechanisms of BF-TK/GCV in GC, MKN-45 cells were treated with PBS, BF-TK, BF/GCV and BF-TK/GCV, respectively. Then the quantitative proteomic profiling was detected by 10-plex TMT. The workflow of TMT-based proteomic analysis was demonstrated in Figure 1A.

**Figure 1.**
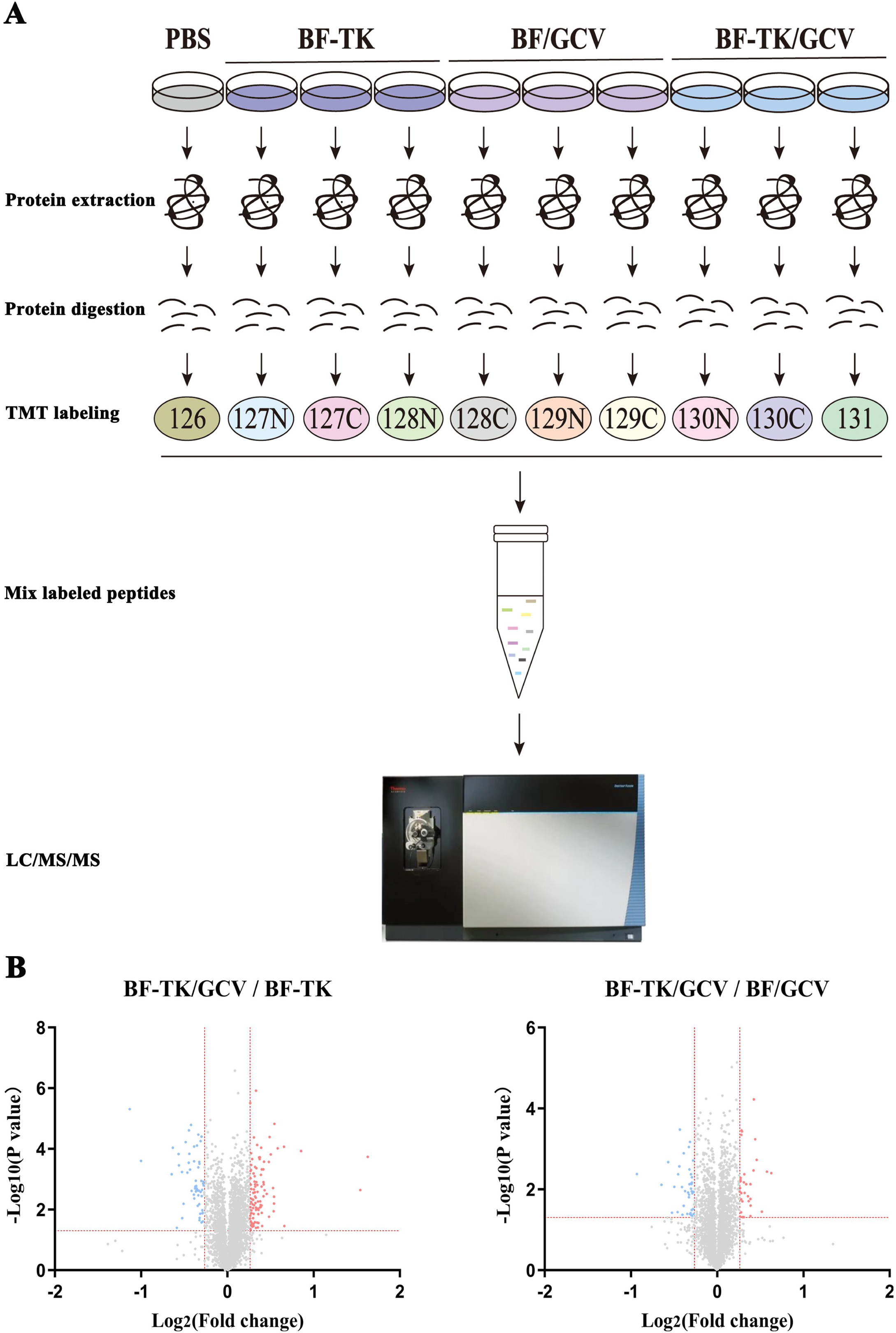
Comparative proteomic analysis of MKN-45 cells treated by BF-TK/GCV. **(A)** Schematic illustration of the TMT-based quantitative proteomic workflow. MKN-45 cells are treated with PBS, BF-TK, BF/GCV or BF-TK/GCV for 48 h and are subjected to 10-plex TMT labeling. Labeled peptides are analyzed by online nano flow liquid chromatography tandem mass spectrometry. **(B)** Identification of differentially expressed proteins in BF-TK/GCV / BF-TK and BF-TK/GCV / BF/GCV groups. Low and high relative expressions are indicated in blue and red, respectively.

Volcano plot filtering was used to identify DEPs with > 1.2-fold changes and p values < 0.05. In the BF-TK/GCV / BF-TK group, 102 proteins were upregulated and 57 proteins were downregulated (Figure. 1B). In the BF-TK/GCV / BF/GCV group, 32 proteins were upregulated and 40 proteins were downregulated (Figure. 1C).

Venn diagrams indicated that 8 and 3 proteins were observed with similar regulation between the BF-TK/GCV / BF-TK and BF-TK/GCV / BF/GCV groups, which suggested that they were important proteins for the effect of BF-TK/GCV (Figure. 2A). Further analysis of the DEPs by unsupervised hierarchical clustering classified these samples into three different cohorts, which also reflected the three distinct characteristics (Figure. 2B).

**Figure 2.**
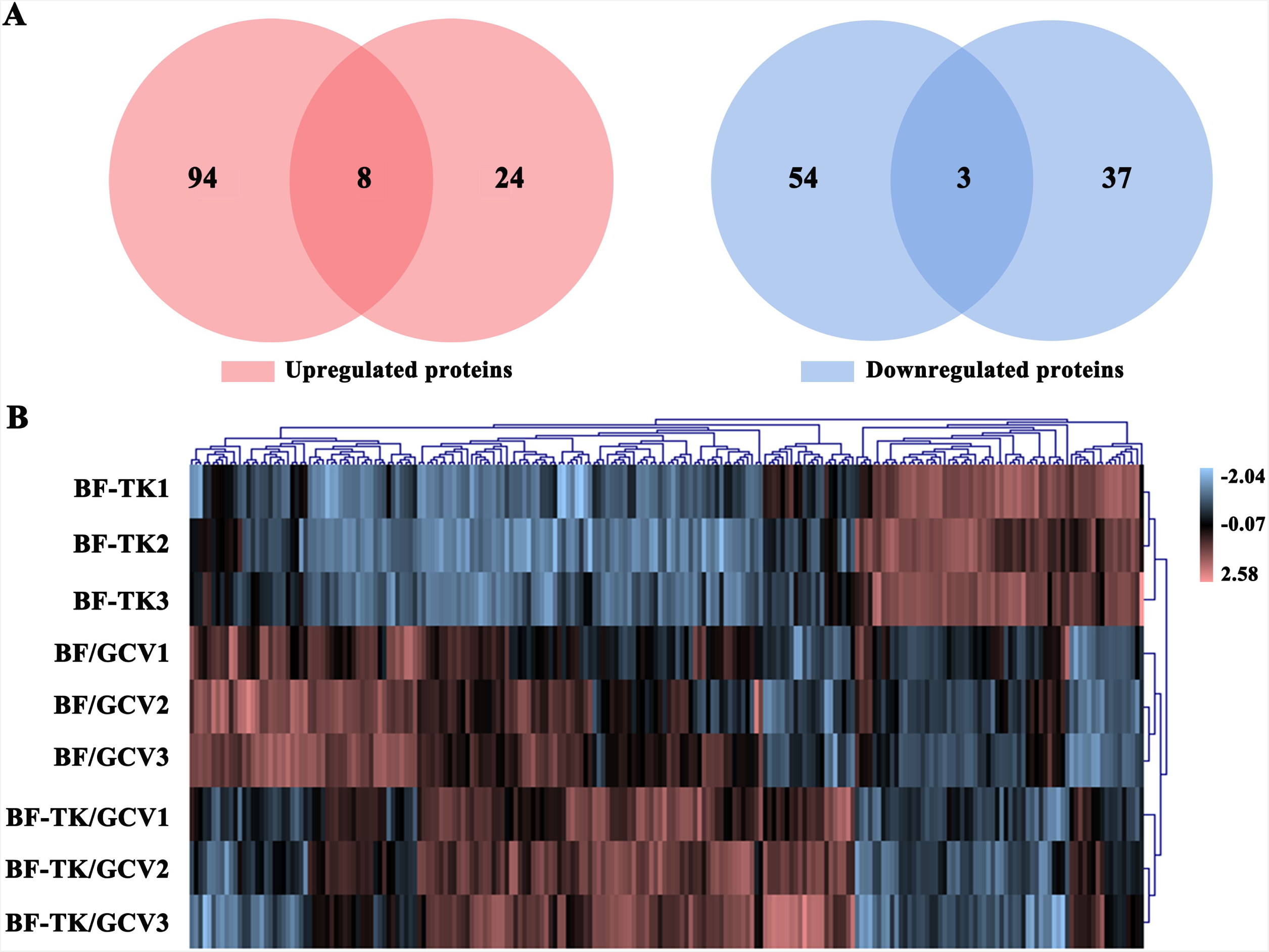
Expression analysis of the identified differentially expressed proteins. **(A)** Venn diagrams display the overlapping differential protein number among the BF-TK/GCV / BF-TK and BF-TK/GCV / BF/GCV groups. **(B)** Clustering of differential proteins from the three groups using a visualized heatmap. Red signifies high relative expression, and blue signifies low relative expression. Various color intensities indicate the expression levels and a log2 scale is used in the color bar.

### Functional enrichment analysis

In order to gain insight into the biological classifications among two groups, the DEPs between groups BF-TK/GCV / BF-TK and BF-TK/GCV / BF/GCV were analyzed separately using the R software based on the GO database. In the BF-TK/GCV / BF-TK group, a total of 2594, 346 and 402 terms in the BP, CC and MF were enriched, respectively. The top 10 enriched BP terms are displayed in Figure 3A. BP analysis revealed significant enrichment for regulation of dendrite development and lysosomal transport. Meanwhile, 1337, 189 and 214 terms were enriched in the BP, CC and MF in the BF-TK/GCV / BF/GCV group. The top 10 enriched BP terms were showed in Figure 3B. BP category analysis indicated that most genes were related to response to nutrient levels and endoplasmic reticulum stress.

**Figure 3.**
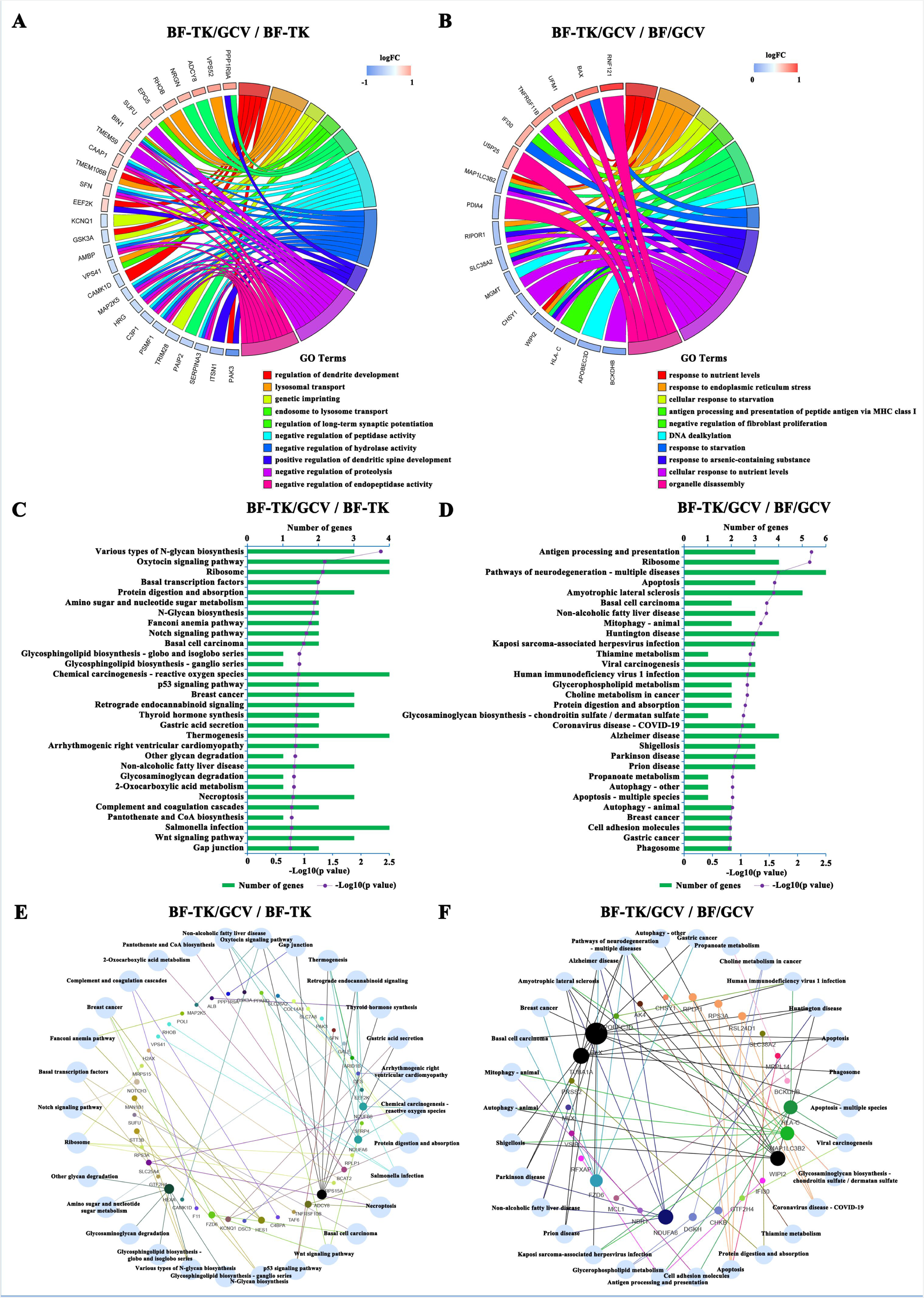
Gene Ontology, Kyoto Encyclopedia of Genes and Genomes pathway enrichment and venn network analysis. Differentially expressed proteins from the **(A)** BF-TK/GCV / BF-TK and **(B)** BF-TK/GCV / BF/GCV groups are analyzed with GO database, and the top 10 enriched BP terms are showed. The top 30 enriched KEGG pathways in the **(C)** BF-TK/GCV / BF-TK and **(D)** BF-TK/GCV / BF/GCV groups are showed. Venn network of top 30 enriched pathways and proteins in the **(E)** BF-TK/GCV / BF-TK and **(F)** BF-TK/GCV / BF/GCV groups.

KEGG pathway enrichment analysis was performed on the two groups (BF-TK/GCV / BF-TK and BF-TK/GCV / BF/GCV) using the R software. The top 30 enriched pathways were showed in Figure 3C and D. In the BF-TK/GCV / BF-TK group, KEGG pathway enrichment revealed that the DEPs were primarily implicated in p53 signaling pathway (TNFRSF10B and SFN) and necroptosis (TNFRSF10B, H2AX and SLC25A4). The KEGG pathway analysis in BF-TK/GCV / BF/GCV group revealed the DEPs mainly participated in apoptosis (BAX, MCL1 and TUBA1A). These pathways associated with cell death which was consistent with our previous studies(Jiang et al, 2013; Jiang et al, 2014; Tang et al, 2009; Wang et al, 2016; Xiao et al, 2014; Yin et al, 2013). In addition, there were pathways such as gap junction (ADCY8 and PPP1R9A) and cell adhesion molecules (VSIR and HLA-C) associated with tumor metastasis. Subsequently, venn network showed the relationships between top 30 enriched pathways and genes in BF-TK/GCV / BF-TK and BF-TK/GCV / BF/GCV groups (Figure. 3E and F).

### PPI networks and module analysis

Further, the PPI networks were constructed based on the STRING database to clarify protein interactions (Figure 4A and B). MCODE was used to find clusters (highly interconnected regions) in the network, and the cluster of the highest score was showed in Figure 4C. In the BF-TK/GCV / BF-TK group, cluster was composed of downregulated proteins, which were AFM, GIG25 (SERPINA3), APCS, HRG, ALB, HPX and AMBP. Meanwhile, MCC method of the CytoHubba plug-in was used to calculate the top 5 proteins as hubproteins. In the BF-TK/GCV / BF-TK group, the top 5 hubproteins were AMBP, HRG, APCS, HPX and GIG25 (Figure 4D), some proteins (HRG, HPX and GIG25) are positively associated with tumor metastasis (Mondaca et al, 2021; Niu et al, 2021; Zhang et al, 2021a). Besides, PPI network showed that CEBPB, NF-κB1, MTOR, HIF-1A, VCAM1 and CXCL12 were interacted directly or indirectly (Figure 4E).

**Figure 4.**
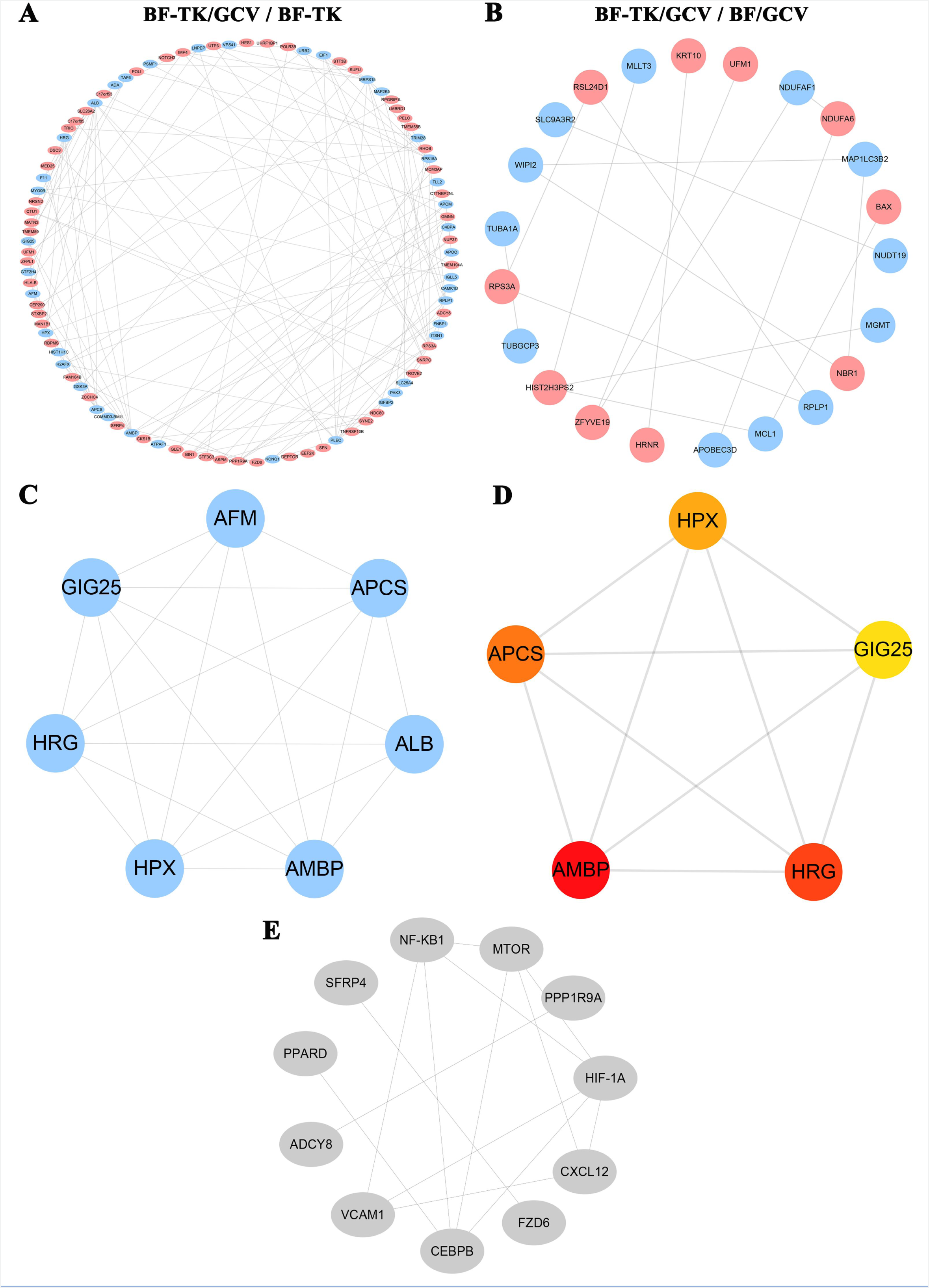
PPI network and subnetwork of the differentially expressed proteins. **(A, B)** Using the STRING online database, the differentially expressed proteins (upregulated proteins are showed in red and downregulated proteins are showed in blue) are filtered into a PPI network complex. The more interactions with other proteins, the more important this protein is. **(C)** Subnetworks are constructed using the MCODE plugin, the subnetwork with highest score are shown. **(D)** Subnetworks are constructed using the CytoHubba plugin, advanced ranking of hubproteins is represented by redder color.

### BF-TK/GCV inhibits gastric cancer metastasis

The gap junction and cell adhesion molecules signaling pathways were related to tumor metastasis. MKN-45 cells treated by BF-TK/GCV showed significantly reduced migration and invasion capabilities as compared to BF-TK or BF/GCV treatment in transwell assays (Figure 5A). Moreover, the six metastasis-related proteins above were further independently validated by the IHC (Figure 5B). The expression of HIF-1A, MTOR, NF-κB1-p105, VCAM1, CEBPB and CXCL12 in BF-TK/GCV group were significantly decreased compared with BF-TK or BF/GCV group (Figure 5C).

**Figure 5.**
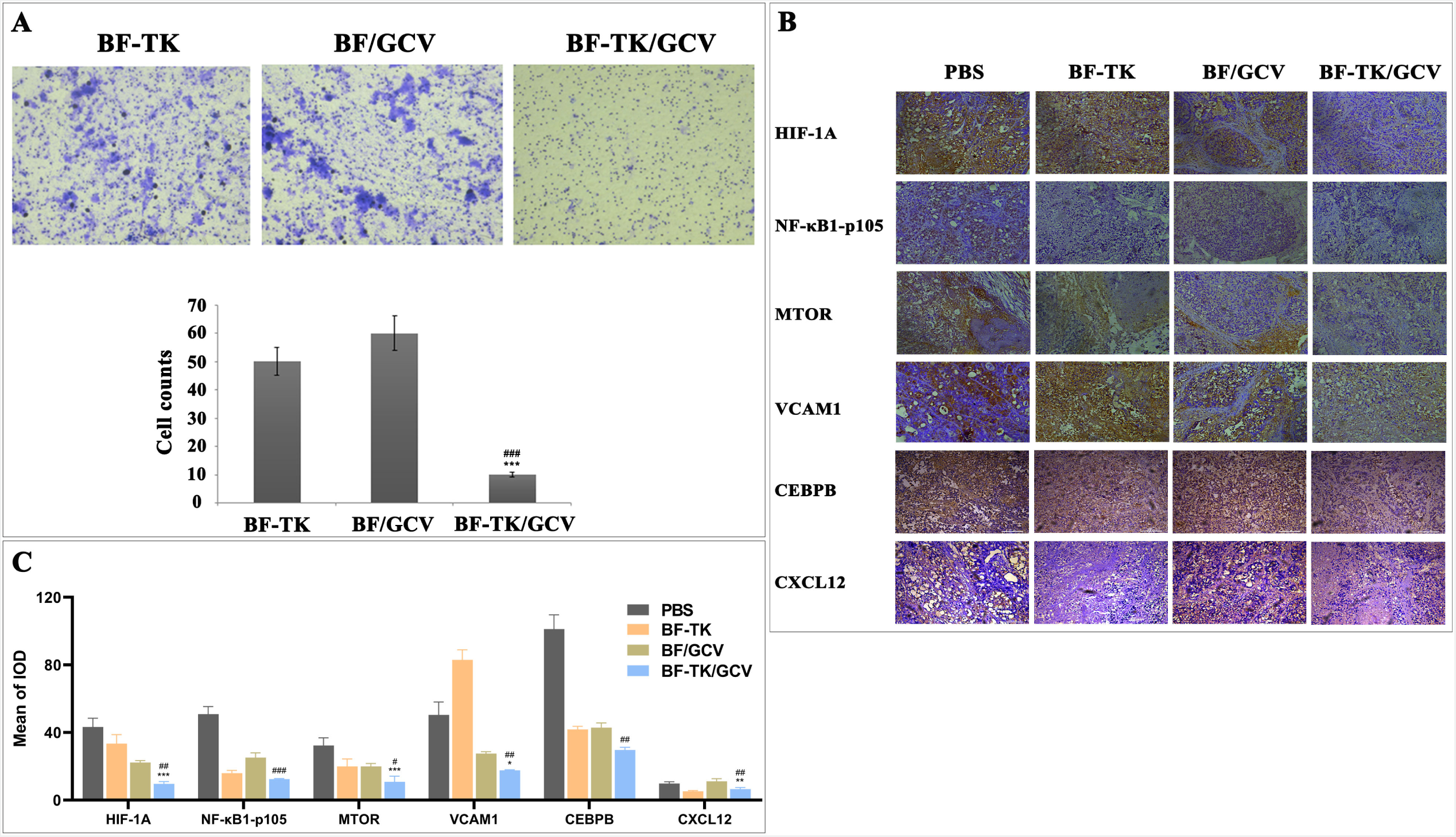
Transwell assay and IHC analysis of metastasis-associated proteins. **(A)** Representative images of transwell assay and quantification of the number of cells which migrate or invade through the basement membrane. **(B)** Representative tissue sections of HIF-1A, MTOR, NF-κB1-p105, VCAM1, CEBPB and CXCL12 in tumor tissues of four groups. **(C)** Quantification of proteins staining; Data represent mean ± SD of three independent experiments. *P<0.05 vs. BF-TK group; **P<0.01 vs. BF-TK group; ***P<0.001 vs. BF-TK group; #P<0.05 vs. BF/GCV group; ##P<0.01 vs. BF/GCV group; ###P<0.001 vs. BF/GCV group.

### Clinical significance of HIF-1A and VCAM1

TCGA data was used to further validate the clinical significance of these proteins. The results revealed that HIF-1A and VCAM1 significantly upregulated in STAD with regional lymph nodes metastasis (N) compared with no regional lymph node metastasis (N0) (Figure. 6A). Moreover, the results suggested that STAD patients with higher expression of VCAM1 exhibited poorer overall survival rate (p=0.026), HIF-1A had a similar trend but not significant, probably because of the smaller number of patients (p=0.064) (Figure. 6B and C). Furthermore, the expression of HIF-1A was positively correlated with that of VCAM1 (R=0.28, p=9.7e-09), suggesting that HIF-1A may play as similar role in the progression of GC as does VCAM1 (Figure. 6D). BF-TK/GCV reduced HIF-1A and VCAM1 expression indicating a good clinical application prospect.

**Figure 6.**
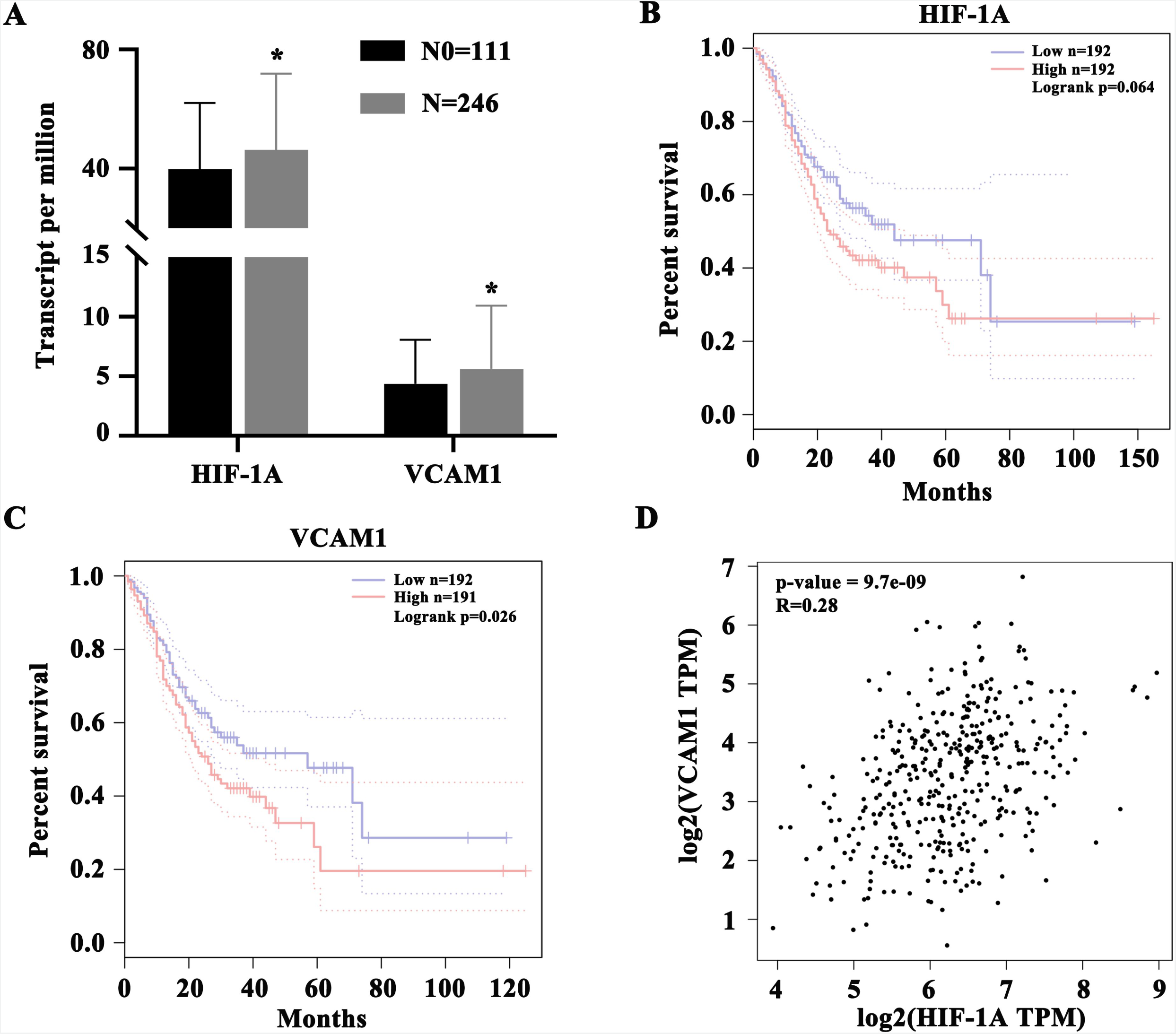
Clinical significance of HIF-1A and VCAM1. **(A)** Expression of HIF-1A and VCAM1 based on nodal metastasis status in TCGA-STAD. **(B, C)** Overall survival analysis of HIF-1A and VCAM1 in TCGA-STAD. **(D)** The correlation between HIF-1A and VCAM1 in TCGA-STAD. Data represent mean ± SD. *P<0.05 vs. N0 group. N0: no regional lymph node metastasis.

## Discussion

Although great progress has been made in cancer treatment over the past decade, effective prevention of tumor metastasis and invasion is still a great challenge for clinical treatment(Ganesh & Massague, 2021; Mazzocca & Carloni, 2009). In this study, we found that BF-TK/GCV inhibits the tumor metastasis, which provided a new strategy for anti-tumor metastasis.

As a model of this study, GC is the fourth leading cause of death and the fifth most common cancer (Sung et al, 2021). The 5-year survival rate of GC patients is still poor because of cancer recurrence due to metastasis (Biagioni et al, 2019). Therefore, the ability of BF-TK/GCV to inhibit GC metastasis provided a new strategy and hope for prolonging the life of GC patients by promoting apoptosis and inhibiting tumor metastasis and invasion.

Delivery vector is crucial to HSV-TK/GCV system. Some viral-mediated HSV-TK/GCV system limited it use for severe immune and inflammatory responses (Hall et al, 1997; Staquicini et al, 2020; Tseng et al, 2016). BF is a nonpathogenic, intrinsic and anaerobic bacterium, which selectively localizes and proliferates in the central area of solid tumors after administrated via the vein or intratumor injection. Based on our previous research that BF-TK/GCV inhibited growth of tumors through multiple mechanisms in our previous studies (Jiang et al, 2013; Jiang et al, 2014; Tang et al, 2009; Wang et al, 2016; Xiao et al, 2014; Yin et al, 2013; Zhou et al, 2016). Here, we further validated that BF-TK/GCV also inhibited tumor metastasis and invasion. We further confirmed that several metastasis-related proteins were decreased expression after BF-TK/GCV treatment, which was consistent with our previous findings that BF-TK/GCV induced metastasis-related VEGF and CD146 downregulation (Jiang et al, 2013; Zhou et al, 2016).

Consistent with our previous findings, this study also proved that several cancer death-associated signaling pathways were enriched in BF-TK/GCV group, including apoptosis, necroptosis and p53 signaling pathways to exert anti-tumor effect (Figure 3C and D) (Jiang et al, 2013; Jiang et al, 2014). Moreover, vacuolar protein sorting (VPS) genes encode a class of proteins involved in vesicular trafficking. VPS52 induces apoptosis via cathepsin D in gastric cancer(Zhang et al, 2017). In this study, VPS41 and VPS52 were upregulated by BF-TK/GCV treatment (Figure 1A).

Moreover, HPX, HRG and GIG25 were significantly downregulated in BF-TK/GCV / BF-TK group. Overexpression of HRG (histidine-rich glycoprotein) inhibits cell proliferation and increased apoptosis in hepatocellular carcinoma(Zou et al, 2020). Hemopexin (HPX) promotes invasion and metastasis of the pancreatic cancer and colorectal carcinoma cells(Niu et al, 2021). GIG25 is a serpin family a member 3 (or SERPINA3) and overexpression of SERPINA3 promotes tumor invasion and migration, epithelial-mesenchymal-transition in triple-negative breast cancer cells(Zhang et al, 2021b). PELO was upregulated by BF-TK/GCV treatment and enriched in mRNA surveillance pathway, However, it was listed out of top 30 pathways (data not showed in Figure 1C). PELO negatively regulates cell migration and metastasis in vivo(Pedersen et al, 2014). The downregulation of these proteins expression suggesting that BF-TK/GCV may inhibit tumor metastasis through these proteins.

There was obviously different with our previous found that gap junction (ADCY8 and PPP1R9A, upregulation) and cell adhesion molecules (VSIR and HLA-C, downregulation) signaling pathways were associated with BF-TK/GCV anti-tumor metastasis.

The low expression of ADCY8 is correlated with poor overall survival and poor progression-free survival in lung adenocarcinoma(Zheng et al, 2021). VSIR (V-set immunoregulatory receptor) is a negative immune checkpoint regulator that inhibits anti-tumor immune responses. Anti-VSIR antibody treatment significantly reduces the number of metastatic nodules in livers of mouse models of PDAC with liver metastases(Hou et al, 2021). It is reported that HLA-C is significant down-regulation in colorectal cancer cells and the cell viability is reduced in the HLA-C overexpression cell line(Lim et al, 2022). BF-TK/GCV was upregulated ADCY8 and PPP1R9A and downregulated of VSIR and HLA-C expression indicated its anti-tumor metastasis mechanism.

IHC assay confirmed several metastasis related proteins were down regulated by BF-TK/GCV treatment (Figure 5C, B). It was reported that reduction of gap junction and HIF1α and CXCR4 expression significantly inhibited the metastasis of breast cancer cells (Zibara et al, 2015). Moreover, neural cell adhesion molecule (NCAM) knockdown inhibits the metastasis of human melanoma cells via the Src/Akt/mTOR/cofilin pathway(Li et al, 2020).

HIF-1 is the major regulator of oxygen homeostasis consists of HIF-1A and HIF-1B subunit. The activation of HIF-1A promotes ovarian cancer cells migration and omental metastasis as well (Sun et al, 2020). In this study, HIF-1A reduced significantly in the BF-TK/GCV group compared with BF-TK or BF/GCV group.

MTOR is a serine/threonine protein kinase as two types of protein complex: mTORC1 and mTORC2. Furthermore, and promotes cancer cell metastasis via miR-451 suppresses glioma cell proliferation and invasion in vitro and in vivo via suppression of the mTOR/HIF-1α/VEGF signaling pathway by targeting CAB39 (Zhang et al, 2019b). In this study, MTOR and HIF-1α were significantly reduced in group BF-TK/GCV compared to BF-TK or BF/GCV (Figure 5).

NF-κB belongs to a family of structurally related eukaryotic transcription factors, the family is divided into p105/p50 (NF-κB1), p100/p52 (NF-κB2), RelA (p65), c-Rel, and RelB. It was reported that NF-κB can positively regulate downstream HIF-1α and VEGF, and associated with the metastasis of ovarian cancer (Yu et al, 2018). BF-TK/GCV increased NF-κB1-p105 (NF-κB1) expression compared to BF/GCV group.

VCAM1 is an important member of the immunoglobulin superfamily and promotes invasion and metastasis of colorectal cancer by inducing transendothelial migration (Zhang et al, 2020). Downregulation of VCAM1 expression blocks breast cancer cell metastasis (Wu et al, 2021). VCAM1 was decreased significantly in the BF-TK/GCV group compared with BF-TK group.

CEBPB is a member of the transcription factor family of CEBP. Study found that STAT3 promoted melanoma metastasis by up-regulating the expression of CEBPB family members (Swoboda et al, 2021). In this study, CEBPB was significantly reduced in group BF-TK/GCV compared to BF-TK or BF/GCV.

CXCL12 is the ligand of CXCR4 and the activation of CXCL12/CXCR4 makes M2 polarized macrophages promote liver cancer metastasis by secreting VEGF (Wang et al, 2020). In this study, CXCL12 was lower in the BF-TK/GCV group compared to BF/GCV group.

Taken together, BF-TK/GCV inhibited GC cell metastasis by downregulating MTOR/HIF-1A pathway and reduced the expression of HIF-1A, MTOR, NF-κB1-p105, VCAM1, CEBPB and CXCL12 confirmed by IHC assay or other metastasis-related proteins (HLA-C, HPX, HRG VSIR and GIG25) confirmed by proteomic assay. It provided a precious data for further studies on BF-TK/GCV antitumor gene therapy. However, the precise functions and roles of these proteins needed to be further explored to better understanding of the underlying mechanisms in the future.

## Material and methods

### Cell and animal treatment

GC cells (MKN-45) were obtained from Committee of Type Culture Collection of Chinese Academy of Sciences (CTCCCAS) and maintained in complete growth medium, RPMI 1640 medium with 10% fetal bovine serum and 1.0%. The cells were cultured in 100-mm culture dishes in a humidified, mixed environment of 37°C and 5% CO_2_. The BF-TK/GCV suicide gene therapeutic system was constructed as our previous describe (Wang et al, 2016; Zhou et al, 2016). MKN-45 cells were treated with PBS, BF-TK, BF/GCV or BF-TK/GCV for 48 h (GCV, 167 µg/ml), respectively.

BALB/c nude mice (male, 3–4 week, 20 g/mouse, n=6) were housed at the Laboratory Animal Center of Chongqing Medical University (Chongqing, China). Mouse model of xenograft tumor was established by injecting MKN-45 cells (1.0 × 10^8^ cells/ml) subcutaneously. Each group was once off directly given PBS, BF-TK, BF/GCV or BF-TK/GCV through intratumor injections (BF or BF-TK was 1.0 × 10^6^ cells/tumor, GCV was 5.0 mg/kg) for 48 h.

This study was carried out in strict accordance with the recommendations in the Guide for the Care and Use of Laboratory Animals of the National Institutes of Health. The protocol was approved by the Committee on the Ethics of Animal Experiments at the Chongqing Medical University (SYXK2012-0001). All surgery was performed under sodium pentobarbital anesthesia, and all efforts were made to minimize suffering.

### Total protein extraction

Cells were suspended in lysis buffer [1.0% sodium deoxycholate (SDS), 8 M urea] with appropriate protease inhibitors to inhibit protease activity. The mixture was allowed to settle at 4°C for 30 min and vortexed at every 5 min, and treated by ultrasound at 40 kHz and 40 W for 2 min. After centrifugation at 16000 g at 4°C for 30 min, the concentration of protein supernatant was determined by bicinchoninic acid (BCA) method by BCA Protein Assay Kit (Pierce, Thermo, USA). Protein quantification was performed according to the kit protocol.

### Liquid chromatography–tandem mass spectrometry (LC-MS/MS) analysis

Protein digestion was performed according to the standard procedure and the resulting peptide mixture was labeled using the 10-plex TMT reagent (Thermo fisher, Art.No.90111) according to the manufacturer’s instructions. Then, the pooled samples were fractionated into fractions by ACQUITY Ultra Performance liquid chromatography (Waters, USA) with ACQUITY UPLC BEH C18 Column (1.7 µm, 2.1 mm × 150 mm, Waters, USA) to increase proteomic depth. At last, labeled peptides were analyzed by online nano flow liquid chromatography tandem mass spectrometry performed on a 9RKFSG2_NCS-3500R system (Thermo, USA) connected to a QExactive Plus quadrupole orbitrap mass spectrometer (Thermo, USA) through a nanoelectrospray ion source.

### Proteomics data analysis

The RAW data were analyzed using Proteome Discoverer (Thermo Scientific, version 2.2) Against uniprot-proteome_UP000005640-Homo sapiens (Human) [9606]-74468s-20190823.fasta database. The MS/MS search criteria were as follows: Mass tolerance of 10 ppm for MS and 0.02 Da for MS/MS tolerance, trypsin as the enzyme with 2 missed cleavage allowed, carbamido methylation of cysteine and the TMT of N-terminus and lysine side chains of peptides as fixed modification, and methionine oxidation as dynamic modifications, respectively. False discovery rate (FDR) of peptide identification was set as FDR ≤ 0.01. A minimum of one unique peptide identification was used to support protein identification. The absolute value of fold change (FC) > 1.20 or < 0.83 and P-value < 0.05 were was adopted as criteria for determining the significance of differential expression of a particular protein (Huang et al, 2020).

### Functional, network and clinical significance analysis

The R (Version 4.1.2, https://www.r-project.org) was used for enrichments of biological processes (BP), molecular functions (MF), and cellular components (CC) based on Gene Ontology (GO) terms. Kyoto encyclopedia of genes and genomes (KEGG) analysis was used to identify enriched pathways (Zhang et al, 2019a). Venn network visualized the relationship between the top 30 enrichment pathways and related genes (Evenn, http://www.ehbio.com/test/venn/) (Chen et al, 2021). Protein–protein interaction (PPI) was assessed using Search Tool for the Retrieval of Interacting Genes (STRING, version 11.5, https,//string-db.org/), and the results under the medium confidence (String score=0.4) were visualized in cytoscape (version 3.9.1, http://www.cytoscape.org/) (Li et al, 2019a; Liao et al, 2021). Moreover, the top 5 hub proteins were identified with MCC method by cytohubba, a plug-in of cytoscape software (Wang et al, 2019). Furthermore, the highest score clusters were analyzed by MCODE plug-in (Duan et al, 2021). The Cancer Genome Atlas-Stomach Adenocarcinoma (TCGA-STAD) RNA-seq gene expression data and clinical data were obtained from the TCGA Data Portal (https://portal.gdc.cancer.gov/) (Xu et al, 2020). The survival probability of high- and low-expression gene groups and gene expression correlation analysis were analyzed by GEPIA (http://gepia.cancer-pku.cn/) (Li et al, 2021b; Xue et al, 2022).

### Transwell assay

For the invasion assay, polycarbonate membrane was coated in the transwell chamber with matrigel (BD Falcon, Bedford, MA, USA). MKN-45 cells (approximately 1×10^5^ in 200 μL serum-free medium) were transferred into the top chamber and added medium with serum in the bottom chamber for 24 h at 37°C. After cells on the top side of the membrane were removed, fixed the cells at the bottom side of the membrane with 4% paraformaldehyde (PFA) and stained with 0.1% crystal violet. Cells at the undersurface of the chamber were imaged and counted.

### Immunohistochemistry (IHC)

IHC of hypoxia inducible factor 1 subunit alpha (HIF-1A), mechanistic target of rapamycin kinase (MTOR), nuclear factor kappa B subunit 1 (NF-κB1)-p105, vascular cell adhesion molecule 1 (VCAM1), CCAAT enhancer binding protein beta (CEBPB) and c-x-c motif chemokine ligand 12 (CXCL12) were detected on MKN-45 tumor xenograft tissues treated by PBS, BF-TK, BF/GCV and BF-TK/GCV, respectively (with three replicates). Retrieved tissues were fixed, decalcified in 10% formalin and embedded in paraffin 24 h posttreatment. The fixed GC tissues of MKN-45 cells were blocked and incubated with HIF-1A (1:200; D122477, Sangon Biotech, China), MTOR antibody (1:200; D160640, Sangon Biotech, China), NF-κB1 (1:200; D120133, Sangon Biotech, China), VCAM1 (1:200; D123530, Sangon Biotech, China), CEBPB (1:200; D155298, Sangon Biotech, China) and CXCL12 (1:200; D161112, Sangon Biotech, China). After being washed, tissues were incubated with biotin-labeled secondary antibody for 30 min, followed by incubation with streptavid in HRP conjugate for 20 min at RT. The presence of the expected protein was visualized by DAB staining and examined under a microscope. Stains with control IgG were used as negative controls.

### Statistical analysis

Statistical analyses were performed with SPSS (Version 25.0, IBM, Chicago, Illinois, USA) software. One-way ANOVA was applied for the comparisons among multiple independent variables. LSD Method was used for multiple pairwise comparisons. Independent sample T test was used for two samples comparisons. The probability level at which difference was considered significant was p < 0.05.

## Abbreviations

TK/GCV: herpes simplex virus thymidine kinase gene with ganciclovir
BF: *Bifidobacterium infantis*
GC: gastric cancer
VEGF: vascular endothelial growth factor
TMTs: tandem mass tags
SDS: sodium deoxycholate
BCA: bicinchoninic acid
FDR: false discovery rate
FC: fold change
BP: biological processes
MF: molecular functions
CC: cellular components
GO: Gene Ontology
KEGG: kyoto encyclopedia of genes and genomes
PPI: protein–protein interaction
STRING: search tool for the retrieval of interacting genes
TCGA-STAD: The Cancer Genome Atlas-Stomach Adenocarcinoma
PFA: paraformaldehyde
IHC: immunohistochemistry
HIF-1A: hypoxia inducible factor 1 subunit alpha
MTOR: mechanistic target of rapamycin kinase
NF-κB1: nuclear factor kappa B subunit 1
VCAM1: vascular cell adhesion molecule 1
CEBPB: CCAAT enhancer binding protein beta
CXCL12: c-x-c motif chemokine ligand 12
DEPs: differentially expressed proteins
STAD: stomach adenocarcinoma
NCAM: neural cell adhesion molecule
ATF: activating transcription factor
MSK1: mitogen- and stress-activated protein kinase 1
TCGA: the cancer genome atlas
HCC: hepatocellular carcinoma;
BMSCs: bone mesenchymal stem cells

## Data availability

All data generated or analyzed during this study are included in this published article and its supplementary files.

## Acknowledgements

This work was supported by grants from the Guangdong Key Laboratory funds of Systems Biology and Synthetic Biology for Urogenital Tumors (2017B030301015) and its Open Grant (2021B030301015-2).

## Authors’ contributions

YM designed the study, CW and JX carried out the experiments, YS undertook the data processing and drafted the original manuscript. YM revised the manuscript and supervised the entire process. All authors contributed to this study and approved the submitted version of the manuscript.

